# Genetic and physiological mechanisms of freezing tolerance in locally adapted populations of a winter annual

**DOI:** 10.1101/640763

**Authors:** Brian J. Sanderson, Sunchung Park, M. Inam Jameel, Joshua C. Kraft, Michael F. Thomashow, Douglas W. Schemske, Christopher G. Oakley

**Author notes:** Department of Genetics, University of Georgia, Athens, GA. Author for correspondence: Christopher G. Oakley.

## Abstract

**Premise of the study:** Despite myriad examples of local adaptation, the phenotypes and genetic variants underlying such adaptive differentiation are seldom known. Recent work on freezing tolerance and local adaptation in ecotypes of *Arabidopsis thaliana* from Sweden and Italy provides the essential foundation for uncovering the genotype-phenotype-fitness map for an adaptive response to a key environmental stress.

**Methods:** Here we examine the consequences of a naturally occurring loss of function (LOF) mutation in an Italian allele of the gene that encodes the transcription factor *CBF2*, which underlies a major freezing tolerance locus. We used four lines with a Swedish genetic background, each containing a LOF *CBF2* allele. Two lines had introgression segments containing of the Italian *CBF2* allele, and two were created using CRISPR-Cas9. We used a growth chamber experiment to quantify freezing tolerance and gene expression both before and after cold acclimation.

**Key results:** Freezing tolerance was greater in the Swedish (72%) compared to the Italian (11%) ecotype, and all four experimental *CBF2* LOF lines had reduced freezing tolerance compared to the Swedish ecotype. Differential expression analyses identified ten genes for which all *CBF2* LOF lines and the IT ecotype showed similar patterns of reduced cold responsive expression compared to the SW ecotype.

**Conclusions:** We identified ten genes that are at least partially regulated by *CBF2* that may contribute to the differences in cold acclimated freezing tolerance between the Italian and Swedish ecotypes. These results provide novel insight into the molecular and physiological mechanisms connecting a naturally occurring sequence polymorphism to an adaptive response to freezing conditions.

## INTRODUCTION

Local adaptation is a consequence of a species occupying heterogeneous habitats. Adaptation to local environments, particularly those where stress imposes strong selection, should involve phenotypic differentiation for ecologically important traits. Such locally adaptive traits are unlikely to be beneficial in contrasting environments (Clausen, Keck, and Hiesey, 1940). More generally, fitness tradeoffs across environments are thought to drive biological diversification at multiple scales (MacArthur, 1972; Futuyma and Moreno, 1988; Whitlock, 1996; Hereford, 2009). Despite many dozens of empirical studies of local adaptation (reviewed in Hereford, 2009), and a growing number of examples mapping the genetic basis of fitness in ancestral environments (Lowry et al., 2009; Hall, Lowry, and Willis, 2010; Ågren et al., 2013; Anderson et al., 2013; Leinonen et al., 2013; Postma and Ågren, 2016; Ågren et al., 2017), there are still very few cases where the traits underlying local adaptation have been identified and experimentally confirmed. Rarer still are examples of the molecular and physiological mechanisms of local adaptation (Anderson, Willis, and Mitchell-Olds, 2011; Savolainen, Lascoux, and Merilä, 2013; Tiffin and Ross-Ibarra, 2014; VanWallendael et al., 2019). Identifying the genes that underlie local adaptation is critical to resolving a longstanding debate about the genetic basis of adaptation (Fisher, 1930; Kimura, 1983; Orr, 1998, 2005; Rockman, 2011; Rausher and Delph, 2015; Remington, 2015; Dittmar et al., 2016). Identifying the traits and physiological mechanisms that underlie local adaptation is a stepping stone to identifying the causal genes, and also informs our understanding of how selection has shaped differentiation in response to environmental heterogeneity.

One trait that is likely to have broad adaptive significance for overwintering species growing at high latitudes or altitudes is freezing tolerance. Freezing tolerance typically requires a period of cold acclimation, an extended period of cold, but non-freezing temperatures (Thomashow, 1999, 2010; Preston and Sandve, 2013; Barrero-Gil and Salinas, 2018), which induces major changes in gene expression, metabolism, and physiology. Typical changes include increased production of soluble sugars and other compounds that decrease the freezing point of the cell, as well as proteins and metabolites to stabilize membranes, reduce or resist ice recrystallization in extracellular spaces, and resist desiccation (Thomashow, 1999, 2010; Preston and Sandve, 2013; Barrero-Gil and Salinas, 2018; Zuther et al., 2018). Thus, cold acclimated freezing tolerance is an example of adaptive phenotypic plasticity, in which cold temperatures trigger an inducible, adaptive physiological mechanism to reduce the stress of freezing temperatures.

Induced responses to stress are thought to reflect adaptations where the mechanism is costly (Agrawal, 2011; Karban, 2011), and where organisms experience stressful environments at different periods in their life history. In this scenario, plants should deploy costly mechanisms of stress tolerance only when the stress is eminent. Unlike induced responses to stress where the cue is closely physically and temporally associated with the agent of selection, such as the induced resistance to herbivores (Agrawal, 2011; Karban, 2011), the cues and the agents of stress in cold acclimated freezing tolerance are temporally separated. In many temperate regions, the conditions that trigger cold acclimation occur far earlier in the life cycle than hard freezing events. The decoupling of cue from the agent of stress in cold acclimation leads to an even greater potential for plants to incur costs. For example, populations in lower latitudes and/or altitudes in the temperate zone are routinely exposed to temperatures that could induce acclimation, yet plants in those environments may never experience severe freezing temperatures. This leads to the hypothesis that response to acclimation cues may be adaptive in some geographic regions where freezing is prevalent and severe, but could result in negative fitness consequences in warmer regions.

Connecting the causal chain between sequence polymorphism, molecular phenotypes, organismal phenotypes, and ultimately fitness in contrasting environments is not an easy task. It is well beyond the scope of any individual study to provide all the necessary information. Detailed studies of the molecular and physiological mechanisms of adaptive traits in study systems for which local adaptation has already been demonstrated in field experiments therefore represent one clear path toward linking sequence polymorphism to ecologically relevant traits and ultimately to fitness. Indeed, in their recent review on stress response networks in plant local adaptation, VanWallendael et al. (2019) highlight that "integrating field-based studies of local adaptation with mechanistic physiological and molecular biology promises advances in multiple areas of plant science.”

Differences in freezing tolerance in locally adapted (Ågren and Schemske, 2012; Ågren et al., 2013; Oakley et al., 2014) ecotypes of *Arabidopsis thaliana* (hereafter *Arabidopsis*) from Sweden (SW) and Italy (IT) represent one such opportunity to uncover the genetic and physiological mechanisms of plant interactions with a stressful environment in the context of local adaptation. Eight years of field experiments have mapped quantitative trait loci (QTL) for local adaptation (Ågren et al., 2013; Postma and Ågren, 2016; Oakley and Ågren, unpublished data). Field and laboratory studies have identified freezing as a major selective agent in SW (Ågren and Schemske, 2012; Ågren et al., 2013; Oakley et al., 2014) and a laboratory study has identified large effect freezing tolerance QTL in the same genomic regions as QTL for local adaptation and fitness tradeoffs (Oakley et al., 2014).

The gene underlying the largest effect freezing tolerance QTL has been identified as encoding the transcription factor *CBF2* (Gehan et al., 2015), and this has been functionally validated using electrolyte leakage freezing tolerance assays on both transgenic and CRISPR mutant lines (Gehan et al., 2015; Park et al., 2018). *CBF2* is well known to be a major regulator of freezing tolerance in both the common laboratory line Col-0, and in natural accessions of *Arabidopsis* (Thomashow, 1999; Alonso-Blanco et al., 2005; Thomashow, 2010; Park et al., 2015; Barrero-Gil and Salinas, 2018), and the IT allele for *CBF2* contains a deletion resulting in a loss of function (Gehan et al., 2015). The CBF genes generally, and *CBF2* in particular, have been shown to mediate large scale changes in gene expression in response to even short-term cold acclimation (Hannah et al., 2006; Gehan et al., 2015; Park et al., 2015; Jia et al., 2016; Zhao et al., 2016; Shi et al., 2017; Park et al., 2018). *CBF2* is therefore a key regulator of adaptive phenotypic plasticity in the SW ecotype of *Arabidopsis* discussed above, and potentially for other winter annuals from freezing climates.

In this study we investigate the molecular mechanisms that underlie the differences in *CBF2* mediated cold acclimated freezing tolerance between our SW and IT ecotypes using a growth chamber freezing assay and RNAseq. We specifically address the effects of the naturally occurring loss of function mutation in *CBF2* in IT by examining gene expression and freezing tolerance of two independent loss of function mutations (produced using CRISPR-Cas9) in the SW genetic background, as well as two near isogenic lines (NILs) where we have introgressed a small part of the Italian genome surrounding *CBF2* into the SW genetic background. We ask the following questions: 1) What proportion of the difference in freezing tolerance between the SW and IT ecotypes can be explained by a *CBF2* loss of function mutation? 2) Do differences in cold responsive gene expression due to the *CBF2* loss of function mutation explain differences in freezing tolerance between the SW and IT ecotypes?

## MATERIALS AND METHODS

### Study system

*Arabidopsis thaliana* is a small selfing (Abbott and Gomes, 1989) annual with a wide native range in Europe, Asia, and Africa (Koornneef, Alonso-Blanco, and Vreugdenhil, 2004; Beck, Schmuths, and Schaal, 2008; Durvasula et al., 2017), where many populations exhibit a winter annual life history (Montesinos et al., 2009; Ågren and Schemske, 2012; Burghardt, Edwards, and Donohue, 2016). In SW, seeds germinate in August and September and seedlings experience low temperatures in the autumn before overwintering as rosettes. In winter in SW, soil temperatures are below freezing for more than 80 days and commonly reach −6°C (Oakley et al., 2014). In IT, seeds germinate in October and November and plants experience cold but non-freezing temperatures throughout the winter as rosettes. Thus, both populations experience temperatures that trigger cold acclimation, but only SW experiences freezing events.

### CRISPR and NIL construction

In order to mimic the loss-of-function mutation in *CBF2* found in IT, we utilized the CRISPR/Cas9 system to generate two independent loss of function *CBF2* mutant lines in the SW genetic background. We followed a multigenerational approach to create the CRISPR lines (Feng et al., 2014). Briefly, the 19 bp oligonucleotides designed to target the coding region of *CBF2* under control of AtU6 promoter were cloned to a single binary vector (pCambia1300): CBF2, 5’-TCGCCGCCATAGCTCTCCG-3’. Seeds generated after floral dip were exposed to antibiotic media to select the first generation of transformed seeds (T1), which were sequenced to confirm the *CBF2* mutation. Transgenic plants with the *CBF2* mutation were self-pollinated for two generations to obtain T3 lines homozygous for the *CBF2* loss-of-function mutations. The T3 lines were then backcrossed to SW in order to remove any possible insertional effects by the T-DNA containing the CRISPR/Cas9 transgene. Two lines were produced (Fig. S1); SW:cbf2 a, which is the same line with a 19 bp deletion in the coding region of *CBF2* in Park et al. (2018), and SW:cbf2 b, with a 13 bp deletion in the coding region of *CBF2*.

We also produced two independent NILs for the *CBF2* region by crossing recombinant inbred lines with IT introgression segments spanning *CBF2* to the SW parental line. The backcrossed lines were then selfed for several generations, and lines of interest were genotyped using a combination of 2b-RAD (Wang et al., 2012) and PCR-based genotyping strategies. Two NILs were ultimately generated and used in experiments: NIL R37, which has a 2.4 Mb introgression segment around the gene *CBF2*, and NIL R38, which has a 6.8 Mb introgression segment that includes *CBF2*. Our use of both CRISPR and NILs in this experiment was motivated by a desire to link these results with field-based estimates of survival and reproduction for plants with functional and non-functional *CBF2* alleles. Due to European Union regulations, lines generated by CRISPR-Cas9 cannot be planted at native field sites, necessitating the use of NILs for the field studies. The inclusion of NILs in addition to the CRISPR lines here allows us to compare the effects of the native LOF allele with those of experimental mutations. Having replicate lines of both types dramatically increases our confidence that the effects we observe in the CRISPR mutants are due to the loss of function of *CBF2* and not to off-target genes.

### Freezing assay

To quantify the effect of the loss-of-function mutation in *CBF2* we exposed seedlings from 6 different lines (IT, SW, the two SW background NILs, and two SW background CRISPRs *CBF2* LOF lines) to a growth chamber freezing assay in which seedlings experienced a period of cold acclimation followed by freezing conditions. The experimental conditions were based on field data, and both this data and protocol have been described previously (Oakley et al., 2014). The experiment was randomized in a stratified fashion in a complete block design. Each block consisted of 2 quartered petri dishes (containing 8 cells), 12 individual seeds of each line were sown in one cell. There were 60 blocks in total, divided evenly among 10 trays (to facilitate randomization within the growth chamber). This entire experimental design was repeated three times, with each temporally separated growth chamber experiment referred to as a batch.

The freezing assay protocol follows Oakley et al. (2014). Briefly, seeds were sterilized using a 30% bleach and TWEEN 20 solution (Sigma Aldrich, St. Louis, Missouri, USA) for 10 minutes and suspended in 0.1% Phytoblend agar (Caisson Laboratories, Inc., Smithfield, Utah, USA) overnight in the dark at 4°C prior to sowing. All seeds were sown on autoclaved Gamborg’s B-5 Basal Salts (without sucrose) and Phytoblend agar (Caisson Laboratories, Inc., Smithfield, Utah, USA) and poured into sterilized petri dishes. The petri dishes were cold stratified in the dark at 4°C for five days to synchronize germination. This was followed by germination and early growth for eight days in a growth chamber at 22°C, 16-hour day length (16L:8D) with a photosynthetically active radiation (PAR) of 125 μmol photons m^−2^ s^−1^. After this period, we put lids on the trays to reduce drying of the agar media and moved the trays to a chamber capable of freezing temperatures to initiate the 10 days cold acclimation phase (4°C, 10L:14D, 50 PAR). We next reduced the temperature to −2°C for 24 hours, and added shaved ice to each cell to facilitate ice nucleation (Smallwood and Bowles, 2002). The chamber then went into freezing at −7 °C for a total of 8 days. During this freezing period the petri dishes were kept in the dark to minimize confounding effects of temperature and photoperiod. To mitigate temperature variation within the chamber, we used supplemental fans and rotated trays twice a day. After the freezing period, we brought the chamber up to 4°C for 24 hours to gradually thaw the plants, followed by 48 hours at 22°C.

We quantified freezing tolerance per cell as mean percent survival after the freezing period. Some cells were not included in the freezing tolerance assay because the plants were sacrificed to collect RNA samples (see below). We excluded seedlings that did not develop true leaves, as preliminary results indicated that seedlings of this size are not freezing tolerant regardless of genotype. Of the total 942 cells included in the freezing assay, we excluded 97 cells because they contained fewer than 4 individual plants of sufficient size to collect freezing tolerance data. In the final dataset, freezing tolerance was estimated for an average of 140.8 cells per line (range = 122-158), each containing an average of 8.26 individual plants — a grand total of 7005 individuals.

Freezing tolerance was analyzed with an analysis of variance with line as a fixed effect. Because of the limited number of batches (3), this factor was treated as a fixed effect. Block nested within batch was treated as a random effect, and significance was tested with a likelihood ratio test. With the exception of IT, which had about five-fold more cell mean freezing tolerance values of zero than the other lines (Fig. S2), the residuals of this model were approximately normally distributed with minimal heteroscedasticity. Reanalysis of a model excluding IT yielded qualitatively similar results for the overall effect of line and the pairwise contrasts to SW (not shown), so we proceed with the full model. Because we are primarily interested in the reduction in freezing tolerance resulting from a non-functional *CBF2* allele, we limited pairwise comparisons to those involving the SW ecotype, and tested these with a-priori linear contrasts. All statistics were performed in JMP v. 13 (JMP, 1989-2019).

### RNA extraction

We randomly selected six blocks in the first batch to be completely harvested for RNA sequencing and these blocks were excluded from the freezing tolerance assay. We harvested all available plant tissue (roots and leaves), four hours after the lights came on in order to minimize the effects of circadian rhythm (Dong, Farre, and Thomashow, 2011). We used RNeasy Plant Mini Kit (Qiagen, Hilden, Germany) for RNA extraction using three replicates of each line at both pre-acclimation (22°C) and post-acclimation (4°C for 10 days) conditions. Total RNA was quantified and checked for quality using a Qubit Fluorometer (Life Technologies Holdings PTE. Ltd., Singapore, Singapore) and a 2100 Bioanalyzer (Agilent Technologies Inc., Santa Clara, California, USA) at the RTSF Genomics Core at Michigan State University.

### Sequencing

Samples were prepared using the Illumina TruSeq Stranded mRNA Library preparation kit (Illumina Inc., San Diego, California, USA) on a Perkin Elmer Sciclone NGS (Perkin Elmer, Inc., Waltham Massachusetts, USA). Completed libraries were quality checked and quantified using a combination of Qubit dsDNA HS (Life Technologies Holdings PTE. Ltd., Singapore, Singapore) and Caliper LabChipGX HS DNA (Perkin Elmer, Inc., Waltham Massachusetts, USA) assays. Libraries were pooled for multiplexed sequencing. Sequencing was carried out in a 1×50bp single end format using Illumina HiSeq 4000 SBS reagents (Illumina Inc., San Diego, California, USA). Base calling was done by Illumina Real Time Analysis (RTA) v2.7.6 and output of RTA was demultiplexed and converted to FastQ format with Illumina Bcl2fastq v2.18.0.

### RNAseq analyses

#### Quality control and mapping

Remaining adapter sequences were removed, bases with quality scores less than 5 were trimmed, and reads smaller than 33bp were excluded using cutadapt v. 1.8.1 (Martin, 2011). Quality of the remaining reads was inspected using FastQC (Andrews, 2010). RNAseq reads from the different genotypes were mapped to the *Arabidopsis thaliana* reference genome (TAIR10) using Tophat version 1.4.1 (Trapnell et al., 2012). TopHat was run in default mode with the following exceptions: the minimum and maximum intron lengths were set to 10 and 15,000 bp, respectively. A GTF file (TAIR10) was used to assist in the mapping of known junctions. Read counts for each gene were obtained using HTSeq 0.6.1 (EMBL, Heidelberg, Germany) using the intersection-noempty option to only include counts for reads mapping to one unique gene.

#### Differential gene expression

Differential expression analysis was implemented in R version 3.0.1 (R, 2011) with the edgeR package v. 3.22.3 (Robinson, McCarthy, and Smyth, 2010). Because estimates of differential gene expression can be inflated by weakly expressed genes, we included only genes with more than one read per million (>1 CPM) in at least two samples. We used the trimmed mean of M-values (TMM) as our normalization method (Robinson, McCarthy, and Smyth, 2010).

Ultimately, we were interested in differential gene expression as an interaction between *CBF2* alleles and the cold acclimation treatment, and the extent to which loss of function mutations in *CBF2* can explain differential expression between IT and SW in response to cold acclimation. The one thing in common among the IT ecotype and the four total CRISPR and NIL lines (in a SW background) is a non-functional *CBF2* allele. We therefore used 5 separate generalized linear models to test for an interaction between genotype and the cold acclimation treatment. SW was included in all five comparisons, and the SW vs. IT comparison establishes ecotypic differences in cold responsive gene expression. Each of the SW vs. CRISPR/NIL comparisons is a measure of the effect of a loss of function mutation in *CBF2* on cold responsive gene expression. Because we have multiple independent comparisons, which is in and of itself an approach to reducing false positives, we took a modified approach to adjusting *P* values for multiple comparisons in order to minimize false negatives. First, we consider only genes where there was a significant (at a Benjamini-Hochberg FDR corrected *P value*, hereafter *P_FDR_*, ≤ 0.05) interaction between genotype and cold acclimation treatment in the SW vs. IT comparison. Then for each of four additional comparisons, we considered only genes that were identified in the comparison between SW and IT (above), and those that had a significant genotype by treatment interaction in their respective pairwise comparison to SW at an uncorrected *P* value of < 0.05. We examined the subset of genes with significant effects of the genotype by cold treatment interaction for expression for all five pairwise comparisons of SW vs. IT, CRISPR, or NIL. These genes were considered to be candidates for downstream targets of *CBF2* that are important in differences in cold acclimated freezing tolerance between SW and IT.

We used RT-qPCR to confirm the results of our RNAseq experiment for two genes that exhibited significant genotype by cold acclimation treatment interactions. RNA was extracted as described above, and RT-qPCR was performed using the Luna Universal Onestep RT-qPCR kit (New England Biolabs, Ipswich, Massachusetts, USA) on a Bio-Rad CFX Connect Real-Time PCR Detection System (Bio-Rad Laboratories Inc., Hercules, California, USA). The threshold quantification cycle (Cq) was determined using Bio-Rad CFX Manager version 3.1. Relative expression ratios were quantified based on the corresponding efficiency of the primers for each gene and the deviation of Cq values for each sample from the mean Cq values of the pre-acclimation samples for each gene (Pfaffl, 2001), in relation to the housekeeping gene *ACT2*.

#### Gene ontology

To assess the function of genes that exhibited significant genotype by environment interactions in the SW vs. IT comparison, we performed a gene ontology (GO) enrichment analysis using the PANTHER v. 14 overrepresentation test, performed with the GO biological processes complete annotations for the complete *Arabidopsis thaliana* gene database (Mi et al., 2019). Fisher exact tests were used to estimate the GO term enrichment *P* values, and a false discovery rate adjustment of *P* values was calculated to correct for the large number of comparisons.

## RESULTS

### Freezing assay

Overall freezing tolerance for the 845 cells (see materials and methods) was 50.2% (SD = 34.5%), and ranged from 0% to 100%. Genotype had a highly significant effect on freezing tolerance (F_5,689_ = 109.5, P < 0.001). This strong signal of genetic-based differences in freezing tolerance accounted for significant variation among batches (F_2,156_ = 32.5, P < 0.001), and significant variation among blocks nested within batch (*X*^2^ = 124.8, df=1, P < 0.001). Least square mean freezing tolerance for SW was 71.9%, which was significantly greater than that of IT of 11.4% (Table 1, Fig. 1). These differences in freezing tolerance between SW and IT are similar to differences in overwinter survival at the Swedish site in cold years (Ågren and Schemske, 2012; Oakley et al., 2014), so differences reported here are reflective of differences in a key fitness component observed in nature. All four lines with a non-functional *CBF2* in the Swedish genetic background had significantly and substantially reduced freezing tolerance compared to SW (Table 1, Fig. 1). Absolute reductions in mean freezing tolerance compared to SW for these 4 lines ranged from 13.11% to 25.74%, explaining from 21.67% to 42.54% of the difference between SW and IT in mean freezing tolerance (Table 1, Fig. 1). Much of the variation among these 4 lines is attributable to the somewhat higher freezing tolerance of NIL R37 compared to the other 3 lines (Table 1, Fig. 1).

**Table 1.** Least square mean freezing tolerance (FrzTol) for each of the six lines in the study. Also given is the reduction in FrzTol of each line compared to SW and results of the linear contrast from the ANOVA testing the significance of this difference. The final column gives the percent of the difference between SW and IT explained by each of the CBF2 loss of function lines in the SW background.

| Line | FrzTol<br>(%) | Reduction compared<br>to SW | Linear contrast<br>compared to SW | Percent difference<br>between SW and IT<br>explained (%) |
| --- | --- | --- | --- | --- |
| SW | 71.9 | n/a | n/a | n/a |
| NIL R37 | 58.8 | 13.1 | $F_{1,688} = 24.8, P < 0.001$ | 21.7 |
| NIL R38 | 50.4 | 21.5 | $F_{1,687} = 66.5, P < 0.001$ | 35.5 |
| SW:cbf2 b | 51.1 | 20.8 | $F_{1,696} = 54.1, P < 0.001$ | 34.4 |
| SW:cbf2 a | 46.2 | 25.7 | $F_{1,695} = 85.7, P < 0.001$ | 42.5 |
| IT | 11.4 | 60.5 | $F_{1,690} = 490.5, P < 0.001$ | n/a |

**Figure 1.**
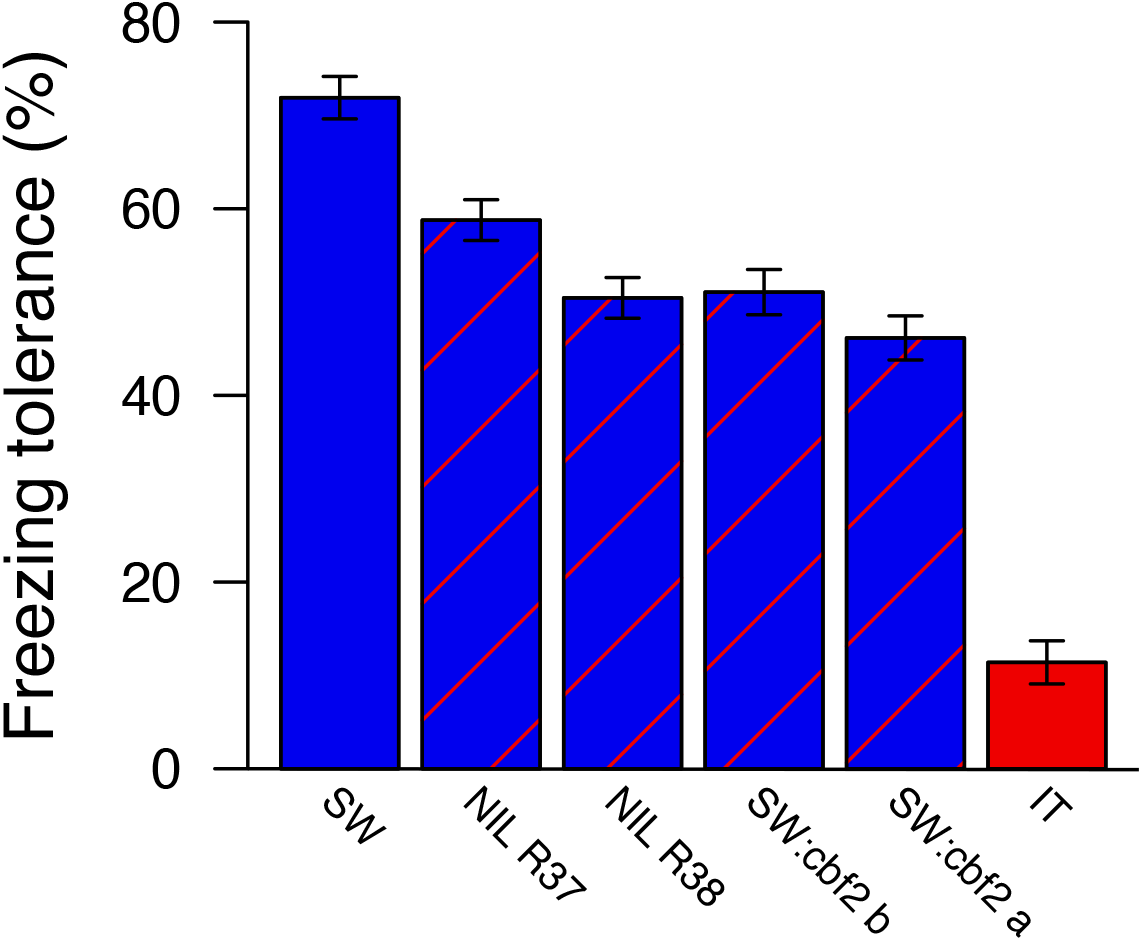
Mean freezing tolerance of SW, IT, and the two NILs and two CRISPR mutant lines containing *CBF2* loss of function alleles in the SW background. Error bars are 1 SE. Linear contrasts comparing SW to each of the other lines were all highly significant (Table 1).

### Gene expression

#### Differences between SW and IT

There were 249 genes that were differentially cold responsive between SW and IT (genotype by treatment interaction) at *P_FDR_* ≤ 0.05 (Fig. S3; Table S1). These genes are involved in genetic pathways involved in glucosinolate biosynthetic processes (6/42 annotated GO terms, *P_FDR_* = 0.00977), response to gibberellin (8/143, *P_FDR_* = 0.0315), alpha-amino acid biosynthetic processes (9/180, *P_FDR_* = 0.03), response to water deprivation (14/346, *P_FDR_* = 0.00665), among others (Table S2).

#### Role of CBF2

Expression of *CBF2* in the warm treatment was very low (less than 0.3 CPM) for all lines, and high for all lines after cold acclimation (range = 17-33 CPM). For the pairwise comparisons of SW to NIL lines, there were 36 and 43 genes for NIL R37 and NIL R38, respectively, that met the above criteria and had a significant genotype by cold acclimation treatment interaction at an uncorrected *P* < 0.05 (Fig. S4, Tables S3 and S4). There were 21 genes meeting both criteria in common between both NILs. For the pairwise comparisons of SW to CRISPR lines, there were 38 and 29 genes for SW:cbf2 a and SW:cbf2 b respectively that met both criteria described above (Fig. S4, Tables S5 and S6). There were 17 genes meeting both criteria in common between both CRISPR lines. There were only 10 cold responsive genes meeting both criteria in common among the 4 NILs and CRISPR lines (Figs 2, 3, S5, and S6).

**Figure 2.**
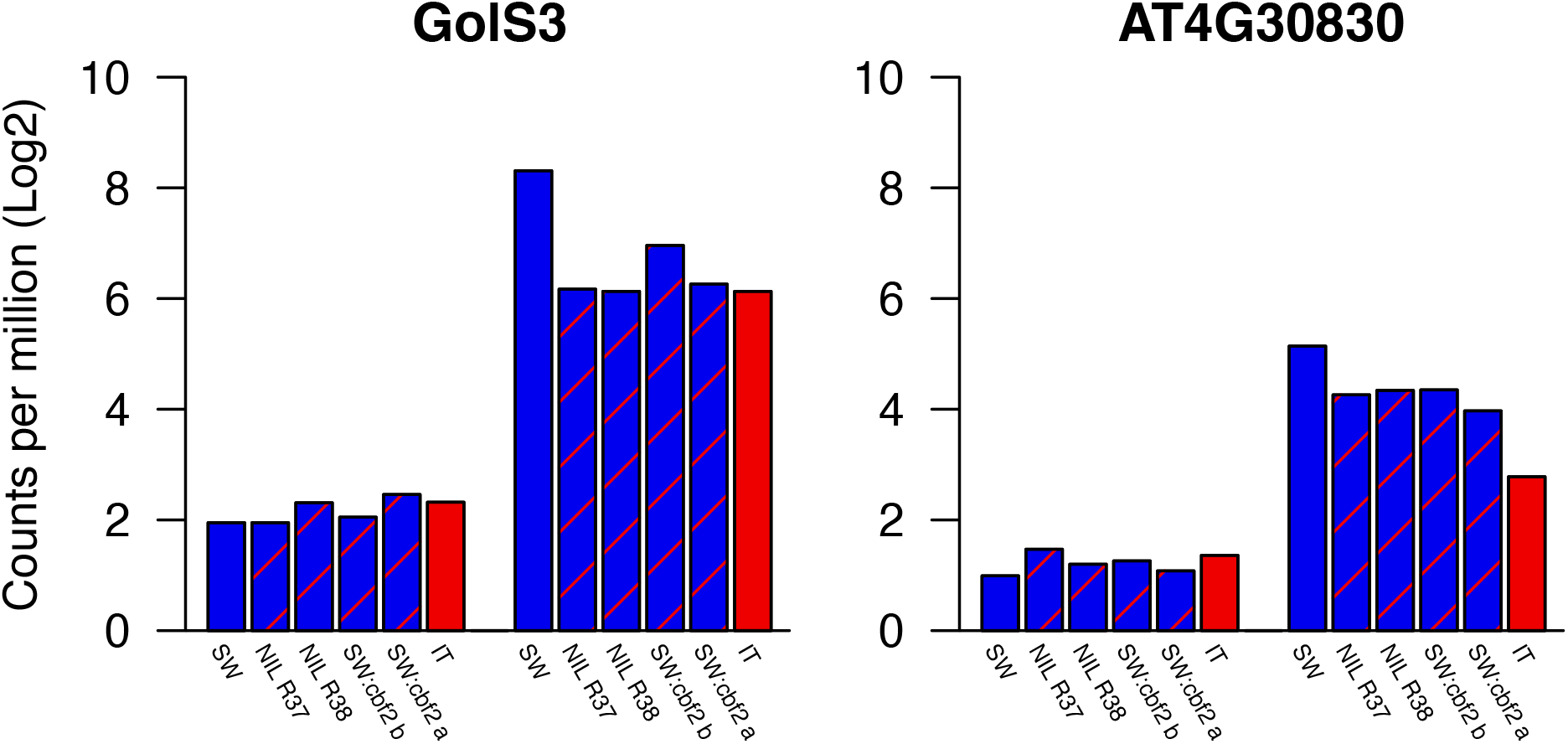
Log_2_ CPM for the most highly cold responsive genes of the 10 candidates before (left group of bars) and after (right group of bars) cold acclimation.

**Figure 3.**
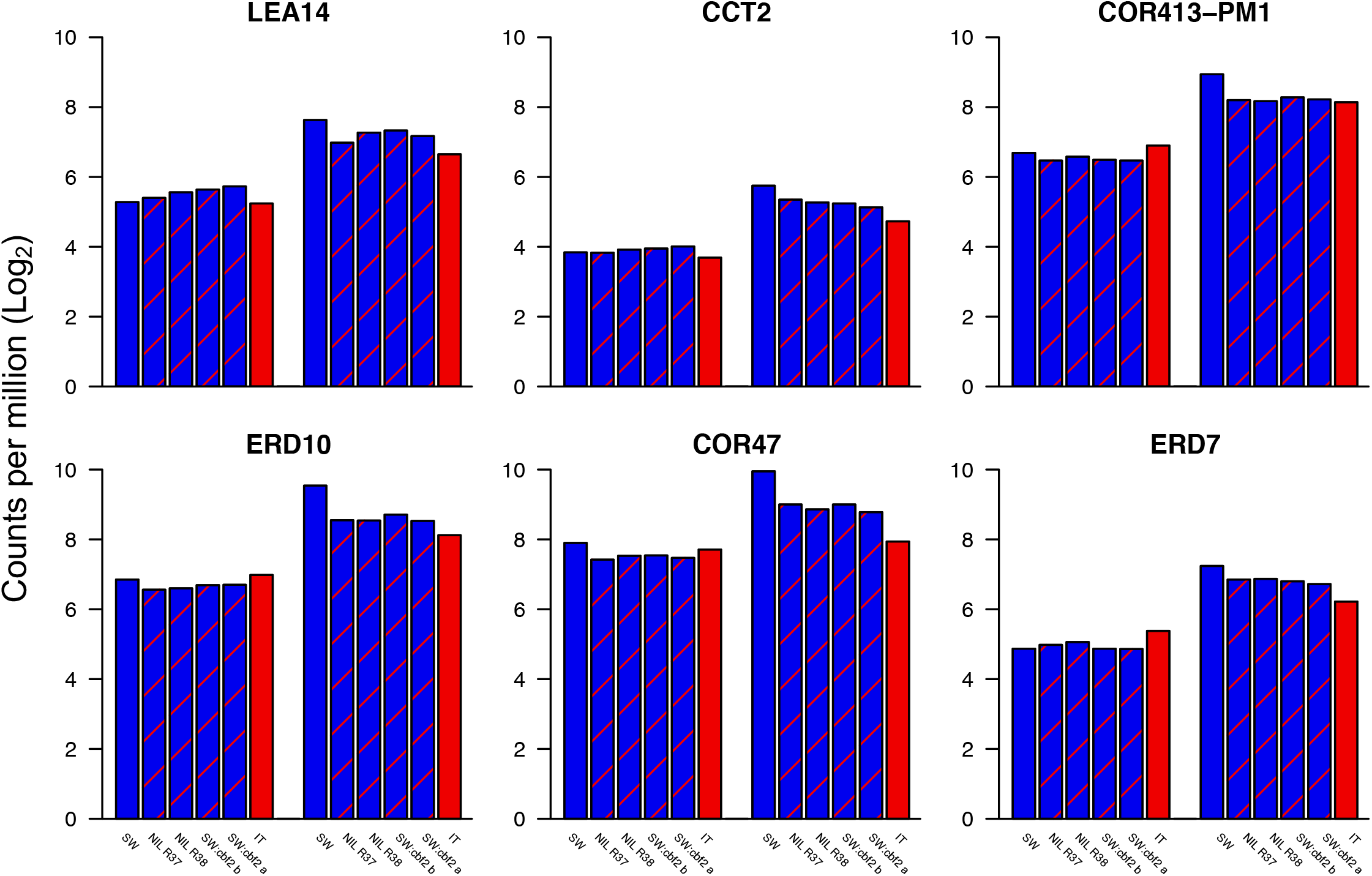
Log_2_ CPM for the remaining highly cold responsive genes of the 10 candidates before (left group of bars) and after (right group of bars) cold acclimation.

To describe patterns of cold acclimated gene expression in SW, we first grouped genes into four categories based on their log2 fold change (LFC) of SW cold acclimation vs. warm (LFC_caSW_; Eq. 1).

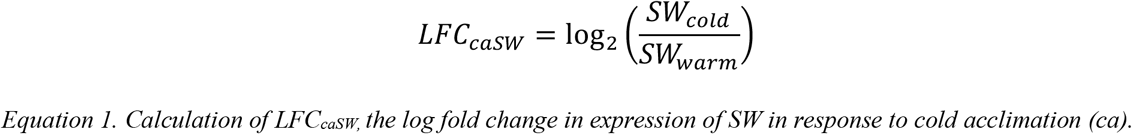

Then, we quantified the difference in cold responsiveness between IT and SW (LFC_caSW_ vs. LFC_caIT_; Eq. 2).

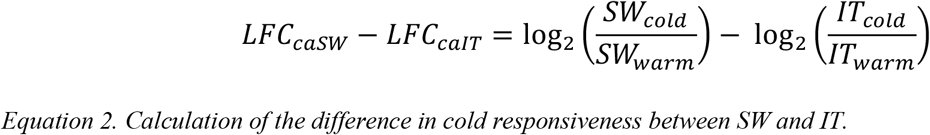

For each of the 4 CRISPR and NIL lines, we calculated the difference in cold responsiveness using Equation 2 substituting the line of interest for IT. Finally, we calculated the average (among the 4 *CBF2* LOF lines in a SW background) proportion of the difference in cold responsiveness between SW and IT (Eq. 2) that can be explained by *CBF2* (Table 2). In other words, we categorized genes first based on how cold responsive they are in SW, then we quantified how much of the difference in cold responsiveness between SW and IT can be explained by LOF mutations in *CBF2*.

**Table 2.** Differential gene expression between pre- and post-cold acclimated plants from SW and IT ecotypes for the 10 genes that were identified as having a significant genotype by treatment interaction (*P_FDR_* ≤ 0.05 for the comparison of IT to SW, and P < 0.05 for all pairwise comparisons between each line and SW). Values are log2 fold change, e.g. a value of 1 represents a 2-fold difference, in expression (*LFC_caSW_* and *LFC_caIT_* as calculated by Equation 1, and *LFC_caSW_* – *LFC_caIT_* as calculated by Equation 2).

| Alias | Gene | $LFC_{caSW}$ | $LFC_{caIT}$ | $LFC_{caSW} - LFC_{caIT}$ |
| --- | --- | --- | --- | --- |
| GolS3 | AT1G09350 | 6.36 | 3.81 | 2.55 |
| NA | AT4G30830 | 4.15 | 1.42 | 2.73 |
| LEA14 | AT1G01470 | 2.35 | 1.42 | 0.94 |
| CCT2 | AT4G15130 | 1.91 | 1.04 | 0.88 |
| COR413-PM1 | AT2G15970 | 2.24 | 1.24 | 1.00 |
| ERD10 | AT1G20450 | 2.70 | 1.14 | 1.56 |
| COR47 | AT1G20440 | 2.05 | 0.23 | 1.82 |
| ERD7 | AT2G17840 | 2.38 | 0.84 | 1.54 |
| NA | AT3G55760 | 1.18 | -0.37 | 1.55 |
| DEAR3 | AT2G23340 | 0.43 | -0.46 | 0.88 |

The first category represents genes that are very highly cold responsive (in terms of log fold change in response to cold) in SW (Figs 2 and S5). We found two genes in this category, *GolS3* (LFC_caSW_ = 6.36) and AT4G30830 (LFC_caSW_ = 4.15). *Gols3* exhibited a striking pattern of cold acclimated gene expression where all 4 lines with LOF mutations in *CBF2* had nearly identical patterns of expression to IT (explaining on average 85% of the log fold difference in cold responsiveness between SW and IT), suggesting that *CBF2* almost completely mediates the difference between SW and IT in cold acclimated gene expression of *GolS3*. For AT4G30830, *CBF2* could explain on average 43% of the log fold difference between SW and IT. The relative expression patterns that we observed using RT-qPCR for GolS3 were consistent with the results we obtained using RNAseq (Fig. S7).

The second category represents highly cold responsive genes (LFC_caSW_ between 1.91 and 2.70) in SW, and included six genes *LEA14, CCT2, COR-413PM1, ERD10, COR47*, and *ERD7* (Figs 3 and S5). Among these, *CBF2* explained the greatest difference in log fold cold responsive gene expression between SW and IT for *LEA14* (80%) and *CCT2* (60%), with lower values for COR-413PM1 (52%), ERD10 (48%), and even lower values for COR47 (35%), and ERD7 (33%). Some of these genes are therefore predominantly regulated by *CBF2*, whereas for others, *CBF2* plays an important, but not predominant role in regulation. The relative expression patterns that we observed using RT-qPCR for COR413-PM1 were consistent with the results we obtained using RNAseq (Fig. S7).

The final two categories of genes are those that are modestly (AT3G55760, LFC_caSW_ = 1.18) or weakly (DEAR3, LFC_caSW_ = 0.43) cold responsive in SW (Figs S5 and S6). Despite the limited cold responsiveness of these genes in SW, *CBF2* could explain a large proportion of differential log fold cold responsiveness between SW and IT, 64% and 82% respectively for AT3G55760 and DEAR3.

## DISCUSSION

Approximately 2/3 of land on earth experiences freezing temperatures at least occasionally during a given year (Larcher, 1980). Freezing tolerance is therefore likely to be a key adaptation to stressful environments for many plants, and because freezing tolerance requires cold acclimation (Thomashow, 1999, 2010; Preston and Sandve, 2013; Barrero-Gil and Salinas, 2018), it is likely to represent adaptive phenotypic plasticity. We investigated the potential genetic and physiological mechanisms of differences in *CBF2* mediated cold acclimated freezing tolerance between locally adapted ecotypes SW and IT. We used CRISPR mutants that mimic a naturally occurring loss of function mutation in *CBF2*, as well as NILs that contain the natural loss of function allele from IT introgressed into an otherwise SW genetic background. For each of these lines we quantified freezing tolerance in a growth chamber experiment, and additionally quantified gene expression before and after cold acclimation. We found that this single mutation in *CBF2* underlies differences in adaptive phenotypic plasticity in the form of cold acclimation between SW and IT, explaining 1/3 of the substantial differential survival through survival between SW and IT. Our approach to identifying the genes that underlie cold acclimated freezing tolerance involved four independent genetic lines, and explicitly tested genotype by treatment interactions for differential gene expression. We were thus able to identify a remarkably short list of ten candidate genes that may play an important role in this adaptive, phenotypically plastic response. Future studies will investigate the contribution of these candidates to local adaptation and fitness tradeoffs using growth chamber and field experiments with NILs that have pairwise combinations of introgressions of *CBF2* and target genes.

### Freezing tolerance

Consistent with previous studies, we find large differences in freezing tolerance between SW and IT. Additionally, the freezing tolerance estimates of the CRISPR and NI lines provide direct evidence for the effect of the naturally occurring LOF mutation in the IT allele of *CBF2* on freezing tolerance. Cold acclimated freezing tolerance of SW was approximately 6.5 fold greater than that of IT (71 vs. 11%, respectively), and these estimates correspond closely to a previous study mapping QTL for freezing tolerance using the same experimental assay (Oakley et al., 2014), and to differences in overwinter survival in the field (Ågren et al., 2013; Oakley et al., 2014). On average, a LOF mutation in *CBF2* in a SW background resulted in a reduction in freezing tolerance of about 20% (Fig. 1) and could explain about 33% of the difference in freezing tolerance between SW and IT. The absolute effect size of 20% observed here is somewhat lower than the 25% estimated for a QTL containing *CBF2* (Oakley et al., 2014), but the percent difference between SW and IT is similar to the 36% from the previous study. The somewhat higher freezing tolerance values for one of the NILs (59% in NIL R37 compared to 50% in NIL R38) is difficult to explain, but the overall pattern for this line follows the expected direction.

Our results add to a growing body of literature pointing to the role of *CBF2* in regulating freezing tolerance, and lays the groundwork for linking the action of *CBF2* with local adaption and fitness tradeoffs across environments. Alonso Blanco *et al*. (2005) were the first to identify and functionally validate the role of a naturally occurring LOF mutation in *CBF2* in cold acclimated freezing tolerance. They identified a large effect QTL for freezing tolerance in a cross between laboratory strain (Ler) and an accession from the Cape Verde Islands (Cvi) that was attributable to a deletion in the *CBF2* allele of Cvi, and confirmed this as the causal variant using transgenic lines in a freezing tolerance assay. In investigating natural variation in freezing tolerance along an elevational gradient in China, Kang *et al*. (2013) also found a mutation in *CBF2* alleles from lower elevation populations that would be predicted to result in LOF. Park et al. (2018) made CRISPR *CBF2* LOF lines in SW, and demonstrated that a LOF mutation in *CBF2* results in increased electrolyte leakage in leaves after freezing. Our work builds upon that of Park et al. (2018) in having an additional independent *CBF2* LOF line, as well as two NILs that can be used directly in future field experiments. Because European Union regulations prevent the planting of lines that carry CRISPR-Cas9 mutations in the field, the only option for connecting patterns of sequence polymorphism to phenotypes to fitness in these contrasting native environments is to combine growth chamber experiments using CRISPR lines and NILs with field experiments using NILs. Furthermore, by quantifying freezing tolerance as survival through freezing, we are able to link this work to extensive long-term field study of the genetic basis of local adaptation in this system.

The discovery of presumably functional variation in CBF genes only in warmer climates (Zhen and Ungerer, 2008; Monroe et al., 2016) is consistent with the idea that cold acclimation is costly in cold, non-freezing environments. Additional indirect support for fitness costs of cold acclimation comes from demonstrations of fitness costs after overexpression of CBF genes (Jackson et al., 2004; Zhen, Dhakal, and Ungerer, 2011). It thus appears that selection on cold acclimation is not merely relaxed in warmer climates (c.f. Zhen and Ungerer, 2008), but rather that the direction of selection changes across environments, such that cold acclimation is favored in some climates and selected against in others. While one direct test of the costs of cold acclimation did not find evidence for such costs (Zhen, Dhakal, and Ungerer, 2011), the short acclimation period used in this experiment may not have been sufficient to induce the full costs of acclimation that might accumulate over long periods in nature. Another confounding factor is that genotypes that have cold acclimated freezing tolerance are also likely to accelerate flowering in response to cold acclimation conditions because these conditions are similar to those promoting vernalization (Preston and Sandve, 2013). The potential confounding effects of vernalization need to be explicitly addressed in future studies on the costs of cold acclimation. We therefore feel that it is premature to conclude, as do VanWallendael et al. (2019), that cold acclimation is cost free in *Arabidopsis*. Our present experiment cannot resolve this question, but fitness data from reciprocal transplant experiments with NILs in SW and IT will directly test the hypothesis of the costs of cold acclimation. In addition, fitness data from growth chamber experiments using CRISPR lines and NILS combined with RNAseq and metabolomic data will allow us to determine the molecular mechanisms of any costs of cold acclimation.

### Gene expression

#### Differences between SW and IT

We identified 249 genes with a significant genotype (SW vs. IT) by cold acclimation treatment interaction (*P_FDR_* ≤ 0.05). A recent study identified 5,200 genes in the SW and IT ecotypes that were differentially expressed in response to cold (Fig. 1D; Gehan et al., 2015). These cold responsive genes include 145 of the genes from our analysis, including nine of the ten most significantly cold-responsive genes (Table S7). The 104 genes from our study that are not included in the Gehan et al. (2015) study include *DEAR3*, which exhibited slight but significant differences in cold-responsive expression in all pair-wise comparisons (See below; Table S8; Fig. S6).

#### Role of CBF2

There has been considerable recent interest in assessing the effects of loss of function CBF mutations on cold acclimated gene expression and freezing tolerance (Zhao et al., 2016; Park et al., 2018). The focus of many of these studies is on the combined effects of loss of function in all three CBF genes to determine what is referred to as the “CBF regulon” for a given accession, rather studying the effects of natural variation in individual CBF genes. Our work builds upon the recent study by Park et al. (2018), who developed CRISPR-Cas9 to mutations in CBF genes in the SW genetic background, with the primary goal of examining freezing tolerance and patterns of cold-acclimated gene expression using RNAseq for a *CBF1, 2*, and *3* triple null mutant. They also reported freezing tolerance for a single *CBF2* null mutant, but did not conduct RNAseq for this line. Here we used the *CBF2* null mutant in Park et al. (2018), an additional independent *CBF2* null mutant, as well as two NILs with IT LOF mutations in a SW genetic background to hone in on the downstream targets of CBF2 that are in part responsible for differences in cold acclimated freezing tolerance, between IT and SW.

Examining the subset of genes with genotype by cold acclimation treatment interactions for all four lines with LOF mutations in *CBF2* in a SW background narrowed the list of differences between SW and IT to just ten genes. Because these genes were identified as significant in all five comparisons of our independent LOF lines (IT, NILs, and CRISPR lines) to SW, we have confidence that these are important candidate genes for cold-acclimated freezing tolerance mediated by *CBF2*. These genes, while not completely regulated by *CBF2*, are likely responsible for most of the differences in freezing tolerance between SW and IT caused by the LOF mutation in the IT *CBF2* allele, and thus are also candidates for mediating the fitness costs of cold acclimation in the Italian environment. The ten genes had annotations with significantly enriched gene ontology terms such as response to abiotic stimulus (GO:0009628), response to stress (GO:0006950), and response to water (GO:0009415; Table S2). We categorize these ten genes based on how cold responsive they are in SW, and further by how much of the difference in cold responsive gene expression between SW and IT can be explained by *CBF2*.

Two of the ten candidates were very highly cold responsive in SW, *GolS3* and AT4G30830. Perhaps the strongest candidate was Galactinol synthase 3 (*GolS3*), which was the most cold responsive gene in SW (Table 2), and for which *CBF2* could explain almost all of the differences in expression between SW and IT. *GolS3* plays a key role in the raffinose biosynthesis pathway, and raffinose is associated with increased tolerance to freezing and other stresses (Taji et al., 2002). *GolS3* has been shown to be cold responsive in a number of studies (Maruyama et al., 2009; Kang et al., 2013; Park et al., 2015; Jia et al., 2016; Zhao et al., 2016), including those using the SW and IT ecotypes (Gehan et al., 2015; Park et al., 2018). Interestingly two studies with triple null mutants of the CBF genes in different genetic backgrounds, Col (Zhao et al., 2016) and SW (Park et al., 2018) both show that the three CBF genes are almost completely responsible for cold acclimated regulation of *GolS3*. Our finding here suggests that *CBF2* explains almost all of the difference between SW and IT in cold responsive gene expression of *GolS3*. Based on our results combined with those of previous studies (Park et al., 2015; Zhao et al., 2016; Park et al., 2018), we therefore conclude that *CBF1* and *CBF3* together regulate the cold responsiveness of *GolS3* that SW and IT have in common.

The other very highly cold responsive gene in SW was AT4G30830, but the expression differences between SW and IT explained by *CBF2* for this gene were more modest (~40%). AT4G30830 is a Myosin-like protein of unknown function (Krishnakumar et al., 2014). This gene has been described as cold responsive in other studies (Gehan et al., 2015; Park et al., 2018). Expression of this gene showed a response similar to *GolS3* in the triple CBF null mutant (Park et al., 2018) suggesting that it is regulated primarily by the CBF genes. Our results taken together with this previous work lead us to conclude that in SW and IT, *CBF1* and *CBF3* may explain the remaining variation in expression differences between SW and IT for AT4G30830. Future studies to identify what, if any, role this gene plays in cold acclimated freezing tolerance would be worthwhile.

The next category of candidates included six genes that were all highly cold responsive in SW: *LEA14*, *CCT2, COR413-PMI, ERD10, COR47*, and *ERD7*. Of these genes, *GolS3, ERD10, COR413-PM1, CCT2*, and AT4G30830 were previously identified as part of the CBF regulon for SW (Park et al., 2018), and the remaining genes have been identified in regulons in other genetic backgrounds (Table S9). The amount of the difference in cold responsive gene expression between SW and IT explained by *CBF2* was variable among this category of genes: *LEA14* (~80%), *CCT2* (~60%), COR413-PMI and ERD10 (~50%), and COR47 and ERD7 (~35%). LEA14 is a Late Embryogenesis Abundant Protein, and is associated with cellular stress, particularly desiccation (Singh et al., 2005). CCT2 is a phosphorylcholine cytidylyltransferase, which acts to increase cellular phosphorylcholine content, an important component of biological membranes, in response to cold (Inatsugi et al., 2009). COR413-PM1 is a multispanning transmembrane protein localized to the plasma membrane that is correlated with freezing tolerance in *Arabidopsis* and cereal crops (Breton et al., 2003), and may play a role in maintaining membrane fluidity under cold temperatures (Su et al., 2018). ERD10 and COR47 are both dehydrin family proteins, thought to play an important role in cellular desiccation resistance, and both have been shown to increase freezing tolerance (Puhakainen et al., 2004). ERD7 is a drought inducible gene that has been shown to be cold responsive (Kimura et al., 2003).

The final two genes in our list of ten candidates were those with only modest or low cold responsiveness in SW. In spite of their relatively small responsiveness on an absolute scale, much of the differences in cold responsiveness between SW and IT for these genes could be attributed to *CBF2* (65% and 80% respectively, for AT3G55760 and DEAR3). Neither of these genes have been previously described as part of the CBF regulon for SW (Park et al., 2018), but both have been identified as part of regulons from other genetic backgrounds (Table S9). AT3G55760 is located in the chloroplast stroma and is involved in starch metabolism (Feike et al., 2016). *DEAR3* is a member of the DREB subfamily ERF/AP2 transcription factors (Sazegari, Niazi, and Ahmadi, 2015), which is the same subfamily of transcription factors as *CBF2*. As a group, the 10 candidates have likely roles in desiccation resistance, sugar biosynthesis or starch metabolism, membrane structure and transport, and regulation of transcription, while some of the functions of these genes are unknown or poorly known.

## CONCLUSIONS

An understanding of local adaptation to stressful environments requires identifying the genetic and physiological changes that confer phenotypic variation, as well as the fitness consequences of such variation in contrasting environments. A comprehensive understanding of the genotype-phenotype-fitness map in nature is beyond the scope of any single study. Detailed molecular studies of traits that have been established as contributing to local adaptation in large multi-year field experiments is perhaps the best approach to connect sequence polymorphism to molecular and organismal phenotypes and ultimately fitness in contrasting environments. The wealth of knowledge about the genetic basis of local adaptation and adaptive traits between Swedish and Italian ecotypes of *Arabidopsis thaliana* (Ågren and Schemske, 2012; Ågren et al., 2013; Oakley et al., 2014; Postma and Ågren, 2016; Ågren et al., 2017; Oakley et al., 2018) provides an excellent foundation from which to pursue the genetic mechanisms of local adaptation. Using a novel approach of examining genotype by environment interactions in gene expression using replicate lines that either simulate (CRISPR) or contain (NILs) the LOF mutation in *CBF2* found in the IT ecotype, we narrowed the list of candidate genes for *CBF2* mediated cold acclimation to just ten genes. These ten genes are excellent candidates for further study of the genetic and physiological changes that underlie the differences in freezing tolerance in these natural populations. Future studies estimating fitness for combinatorial NILs containing pairwise combinations of IT alleles of *CBF2* and each of these ten genes in growth chambers and in the field will be used to investigate interactions between *CBF2* and downstream targets. Such experiments will be coupled with metabolite analysis, particularly steps related to raffinose biosynthesis, to provide additional insight into the mechanisms underlying adaptive cold acclimation responses, and the potential for these mechanisms to result in fitness costs in alternate environments.

## ACKNOWLEDGEMENTS

The authors would like to thank A. Babbit, J. Eilers, and M. Kargul, who helped to implement the freezing tolerance assays, and the RTSF Genomics Core at MSU for sequencing and assistance. We are grateful to Dr. Jian-Kang Zhu, Purdue University, for providing the plasmid used to generate the CRISPR lines. We thank A. Berardi, R. Deater, N. Mano, R. Watson, and members of the Zhang Lab at Purdue for assistance with qPCR. We thank members of the Aime, McAdam, McNickle, and Mickelbart labs at Purdue for helpful comments on a draft of the manuscript. Funding was provided by NSF DEB grant (1743273) to CGO, DWS, and MFT, and by AgBioResearch at Michigan State University to MFT.

## AUTHOR CONTRIBUTIONS

C.G.O., D.W.S, and M.F.T. conceived of the study, C.G.O. and S.P. designed the experiment, S.P. and C.G.O. developed the CRISPR and NIL lines respectively, M.I.J., S.P., J.C.K, and C.G.O. carried out the experiment, C.G.O., B.J.S, and S.P. analyzed the data and produced the figures, B.J.S. and C.G.O. drafted the manuscript with help from S.P. and M.I.J., and all authors contributed to revising the manuscript.

## DATA ACCESSIBILITY STATEMENT

Upon acceptance, all data will be deposited in the Dryad data depository and/or other appropriate publicly available data repositories.

## SUPPLEMENTARY MATERIAL

**Table S1.** List of genes with significant gene by environment interactions in our study for the pairwise comparison of IT to SW (*P_FDR_* ≤ 0.05). F, PValue, and FDR refer to the significance test of the interaction term.

**Table S2.** Gene ontology enrichment for the 249 genes with significant genotype by environment interactions for the pairwise comparison of IT to SW (*P_FDR_* ≤ 0.05).

**Table S3.** List of genes with significant differences in cold responsive expression between SW and NIL R37 (*P* value < 0.05), and which also in Table S1. F, PValue, and FDR refer to the significance test of the interaction term.

**Table S4.** List of genes with significant differences in cold responsive expression between SW and NIL R38 (*P* value < 0.05), and which also in Table S1. F, PValue, and FDR refer to the significance test of the interaction term.

**Table S5.** List of genes with significant differences in cold responsive expression between SW and CRISPR line cbf2 a (*P* value < 0.05), and which also in Table S1. F, PValue, and FDR refer to the significance test of the interaction term.

**Table S6.** List of genes with significant differences in cold responsive expression between SW and CRISPR line cbf2 b (*P* value < 0.05), and which also in Table S1. F, PValue, and FDR refer to the significance test of the interaction term.

**Table S7.** List of genes with significant gene by environment interactions in our study for the pairwise comparison of IT to SW (*P_FDR_* ≤ 0.05) that are included among those identified by Gehan et al. (2015).

**Table S8.** List of genes with significant gene by environment interactions in our study for the pairwise comparison of IT to SW (*P_FDR_* ≤ 0.05), which are not present in Gehan et al. (2015).

**Table S9.** Comparison of genes included in Figure S5 with previously published CBF regulons from four genetic backgrounds. SW and IT from Park et al. (2018), Col-0 from Zhao et al. (2016), and WS from Park et al. (2015).

**Figure S1.** Three transcription factor encoding CBF genes in tandem array in the SW and IT ecotypes, and two lines with CRISPR-induced mutations in the *CBF2* gene. Open boxes indicate *CBF1, CBF2*, and *CBF3* coding regions (shaded portion indicates DNA binding domain) in the order they are arranged in the genome. The green lines indicate the transcription activation domains. The filled triangles indicate site of the naturally occurring 13 bp deletion in the IT *CBF2* gene. Open triangles indicate the two independent CRISPR induced deletion. SW:cbf2 a is the same as in Park et al. (2018).

**Figure S2.** Distribution of mean freezing tolerance values per cell. The distribution of values for all six lines combined are given in black, and the distribution of values for just IT are given in red.

**Figure S3.** Heat map of expression differences for the 249 genes that were identified as having a significant genotype by treatment interaction (*P_FDR_* ≤ 0.05 for the comparison of IT to SW). The value of each cell represents the log2 transformed fold-change in gene expression, calculated as the quotient of normalized counts-per-million averages for each line between cold and warm treatments. Yellow-red color represents genes that are highly expressed in the cold treatment, while black-purple color represents genes that are highly expressed in the warm treatment. Plot generated using the heatmap.2 function in the R package plots (Warnes et al., 2019).

**Figure S4.** Venn diagram of genes with a significant genotype by treatment interaction (*P_FDR_* ≤ 0.05) between IT and SW, and P < 0.05 for each pairwise comparison of the NILs and CRISPR lines to SW. Plot generated using the venn.diagram function in the R package VennDiagram (Chen, 2018).

**Figure S5.** Heat map of expression differences for the 10 genes that were identified as having a significant genotype by treatment interaction (*P_FDR_* ≤ 0.05 for the comparison of IT to SW, and P < 0.05 for all pairwise comparisons between each line and SW). The value of each cell represents the log2 transformed fold-change in gene expression, calculated as the quotient of normalized counts-per-million averages for each line between cold and warm treatments. Yellow-red color represents genes that are highly expressed in the cold treatment, while black-purple color represents genes that are highly expressed in the warm treatment. Plot generated using the heatmap.2 function in the R package plots (Warnes et al., 2019).

**Figure S6.** Log_2_ CPM for the least responsive genes of the 10 candidates before (left group of bars) and after (right group of bars) cold acclimation.

**Figure S7.** Relative expression for *GolS3* and *COR413-PM1* quantified by RT-qPCR, normalized to expression of the housekeeping gene *ACT2*. Each biological replicate was run in triplicate for three technical replicates. Points are means of three biological replicates, and error bars are the standard error of those means. Primer sequences used are as follows: *GolS3* F: 5-TGTGCCAAAGCTCCATCCGC-3, *GolS3* R: 5-TGGTGTTGACAAGAACCTCGCT-3, *COR413-PM1* F: 5-TGCTGGCACATTCAGAGACAG-3, *COR413-PM1* R: 5-CAGACGGGGAAGACGACGAGA-3, *ACT2* F: 5-CTGGATCGGTGGTTCCATTC-3, *ACT2* R: 5-CCTGGACCTGCCTCATCATAC-3.

