## Supplemental Figures for "Genetic and physiological mechanisms of freezing tolerance in locally adapted populations of a winter annual"

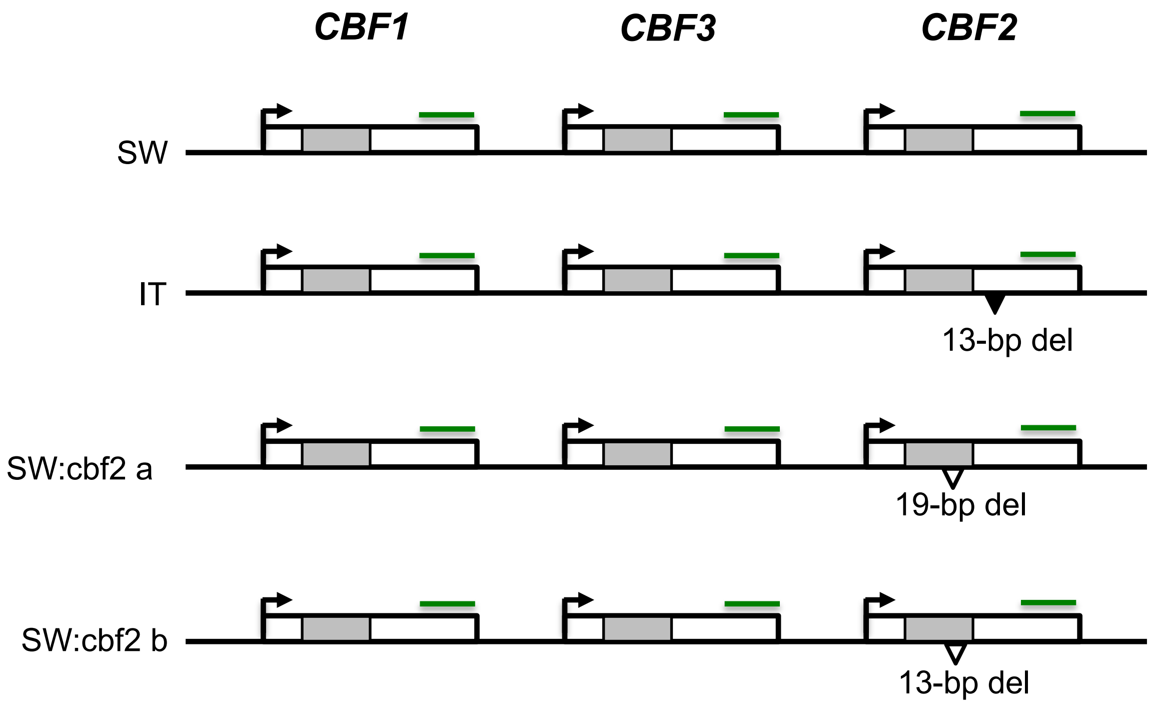


**Figure S1.** Three transcription factor encoding CBF genes in tandem array in the SW and IT ecotypes, and two lines with CRISPR-induced mutations in the *CBF2* gene. Open boxes indicate *CBF1*, *CBF2*, and *CBF3* coding regions (shaded portion indicates DNA binding domain) in the order they are arranged in the genome. The green lines indicate the transcription activation domains. The filled triangles indicate site of the naturally occurring 13 bp deletion in the IT *CBF2* gene. Open triangles indicate the two independent CRISPR induced deletion. SW:cbf2 a is the same as in Park et al. (2018).

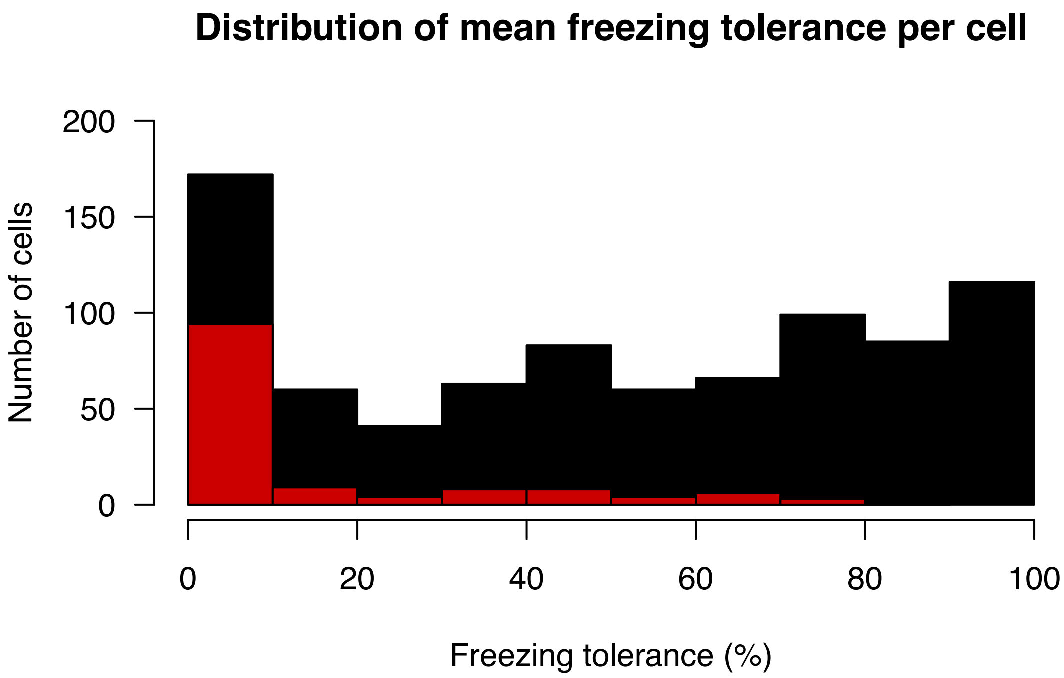


**Figure S2.** Distribution of mean freezing tolerance values per cell. The distribution of values for all six lines combined are given in black, and the distribution of values for just IT are given in red.

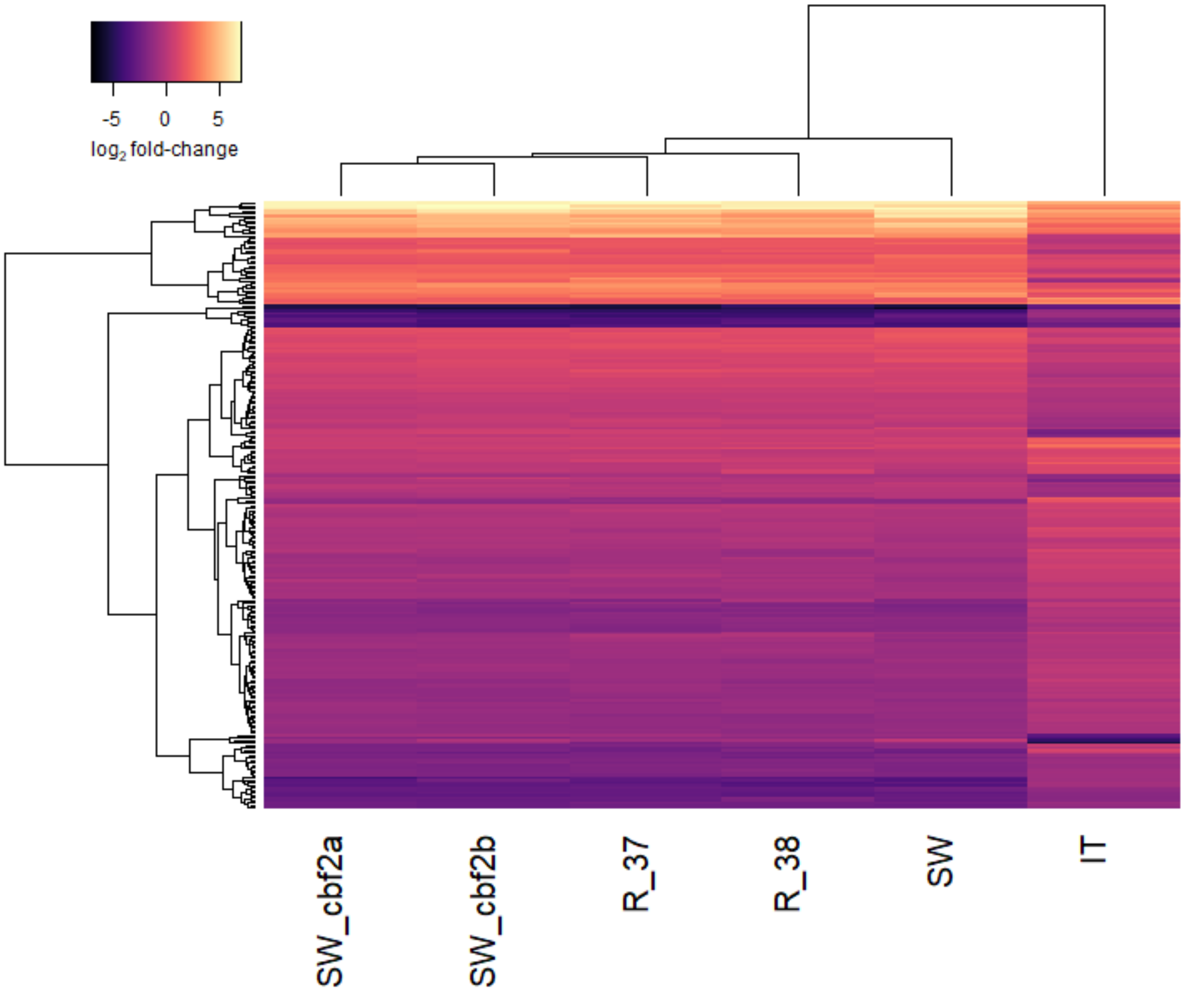


**Figure S3.** Heat map of expression differences for the 249 genes that were identified as having a significant genotype by treatment interaction ($P_{FDR} \leq 0.05$ for the comparison of IT to SW). The value of each cell represents the $\log_2$ transformed fold-change in gene expression, calculated as the quotient of normalized counts-per-million averages for each line between cold and warm treatments. Yellow-red color represents genes that are highly expressed in the cold treatment, while black-purple color represents genes that are highly expressed in the warm treatment. Plot generated using the heatmap.2 function in the R package plots (Warnes et al., 2019).

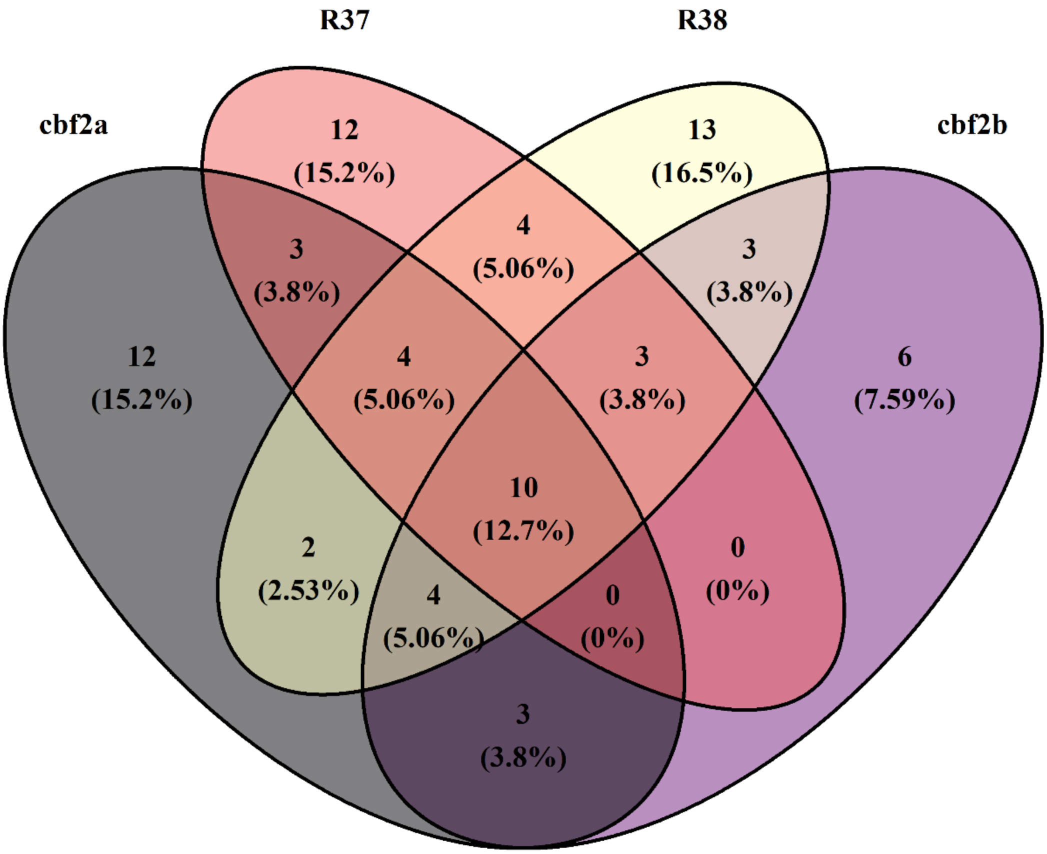


**Figure S4.** Venn diagram of genes with a significant genotype by treatment interaction ($P_{FDR} \leq 0.05$) between IT and SW, and P < 0.05 for each pairwise comparison of the NILs and CRISPR lines to SW. Plot generated using the venn.diagram function in the R package VennDiagram (Chen, 2018).

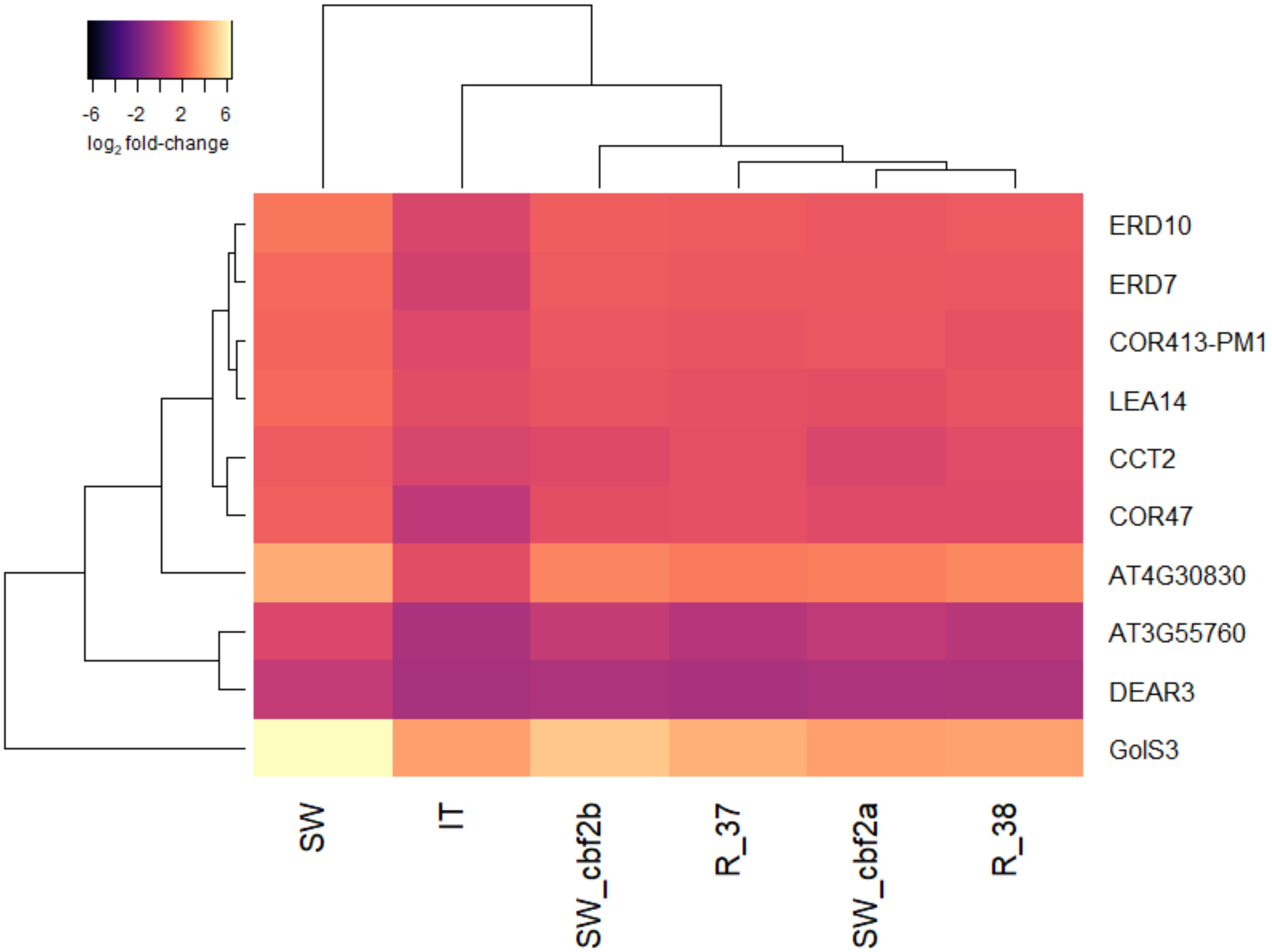


**Figure S5.** Heat map of expression differences for the 10 genes that were identified as having a significant genotype by treatment interaction ($P_{FDR} \leq 0.05$ for the comparison of IT to SW, and $P < 0.05$ for all pairwise comparisons between each line and SW). The value of each cell represents the $\log_2$ transformed fold-change in gene expression, calculated as the quotient of normalized counts-per-million averages for each line between cold and warm treatments. Yellow-red color represents genes that are highly expressed in the cold treatment, while black-purple color represents genes that are highly expressed in the warm treatment. Plot generated using the heatmap.2 function in the R package plots (Warnes et al., 2019).

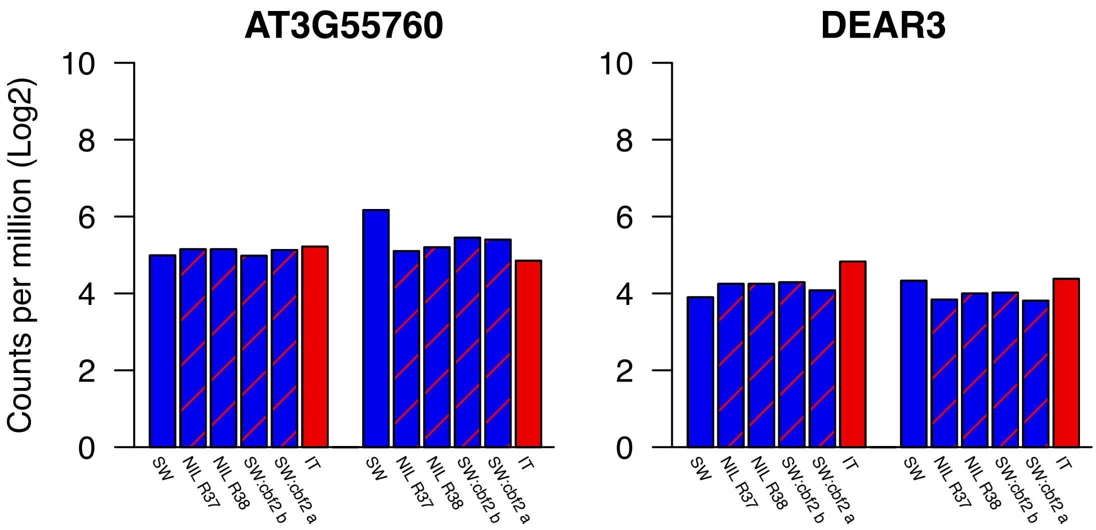


**Figure S6.** $Log_2$ CPM for the least responsive genes of the 10 candidates before (left group of bars) and after (right group of bars) cold acclimation.

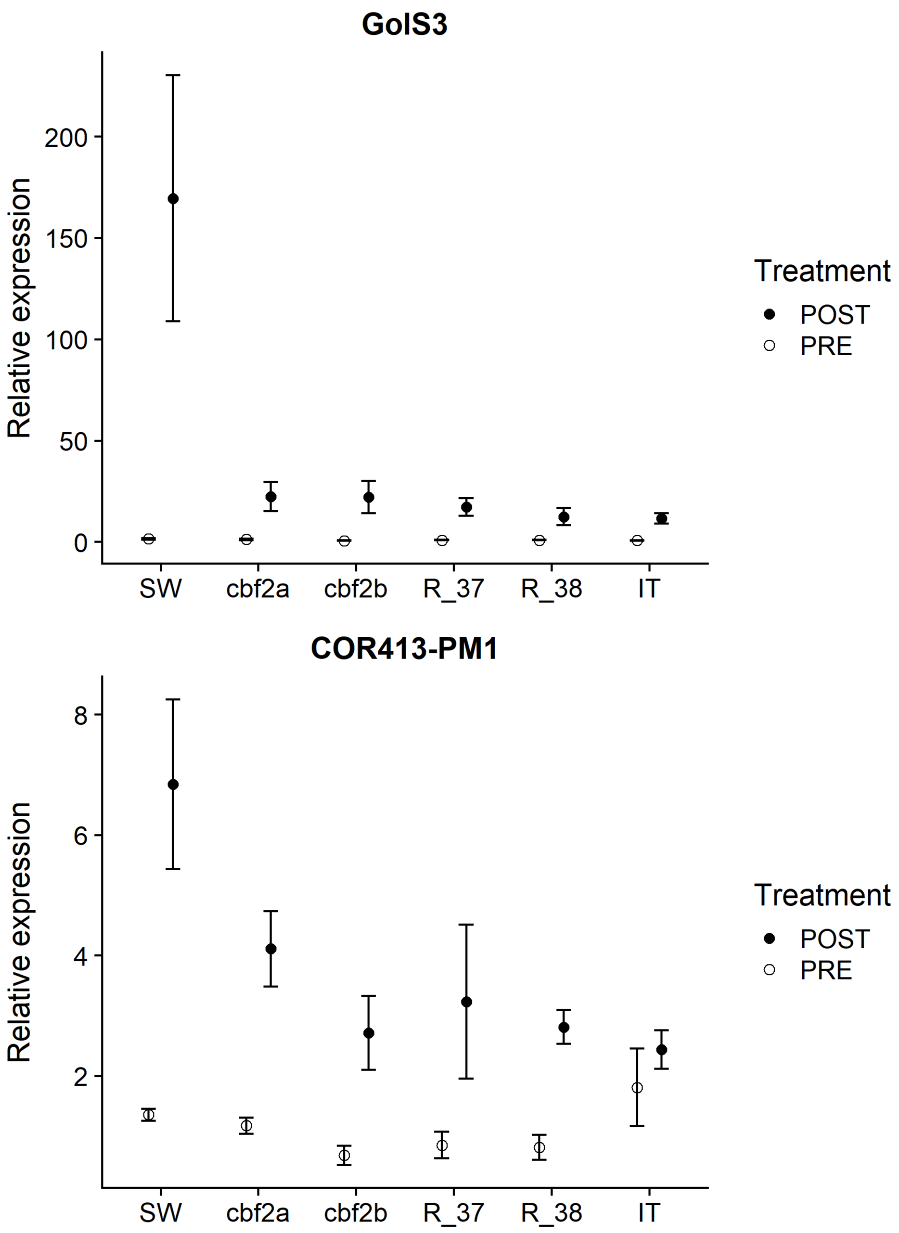


**Figure S7.** Relative expression for *GolS3* and *COR413-PM1* quantified by RT-qPCR, normalized to expression of the housekeeping gene *ACT2*. Each biological replicate was run in triplicate for three technical replicates. Points are means of three biological replicates, and error bars are the standard error of those means. Primer sequences used are as follows: *GolS3* F: 5-TGTGCCAAAGCTCCATCCGC-3, *GolS3* R: 5-TGGTGTTGACAAGAACCTCGCT-3, *COR413-PM1* F: 5-TGCTGGCACATTCAGAGACAG-3, *COR413-PM1* R: 5-CAGACGGGGAAGACGACGAGA-3, *ACT2* F: 5-CTGGATCGGTGGTTCCATTC-3, *ACT2* R: 5-CCTGGACCTGCCTCATCATAC-3.
